# Intranasal CRISPR- lipid nanoparticles targeting MAPK9 reduce neuroinflammation after traumatic brain injury

**DOI:** 10.64898/2026.04.25.720847

**Authors:** Goknur Kara, Yaqoob Ali, Jessica López-Espinosa, Peter Park, Morgan Holcomb, Hannah Flinn, Noah Taylor, Tyler Galbraith, Fransisca Leonard, Sonia Villapol

**Author notes:** **Corresponding Authors:** Sonia Villapol, Ph.D., Fransisca Leonard, PhD.

## Abstract

Traumatic brain injury (TBI) triggers a sustained neuroinflammatory response driven by activated microglia, which contributes to secondary injury and long-term neurological dysfunction. Therapeutic reprogramming of microglial activation from a pro-inflammatory (M1-like) to a reparative (M2-like) phenotype represents a promising strategy; however, the lack of cell-specific targeting within an injured brain has limited clinical translation. Here, we developed a targeted gene-editing nanotherapy to modulate post-traumatic innate immune responses. Lipid nanoparticles (LNPs) encapsulating CRISPR-Cas12a components were engineered to target mitogen-activated protein kinase-9 (MAPK9), a key regulator of pro-inflammatory signaling, and were conjugated with an Iba-1 antibody (Iba-1-CRISPR-LNPs) to enable selective targeting of microglia. *In vitro*, MAPK9 editing in primary macrophages inhibited M1 polarization and promoted an M2-like phenotype, leading to reduced production of pro-inflammatory cytokines. In a TBI mouse model, intranasal administration of Iba-1-CRISPR-LNPs achieved efficient delivery to the injured brain, with selective localization in Iba-1+ microglia. MAPK9 CRISPR targeting significantly attenuated microglial activation, reduced central and peripheral inflammatory responses, and decreased pro-inflammatory cytokine levels. Importantly, this approach demonstrated a favorable safety profile, with no detectable toxicity across major organs. Collectively, these findings establish a non-viral, intranasal CRISPR-based strategy for cell-specific modulation of neuroinflammation following TBI. Targeted genome editing of MAPK9 effectively reprograms microglial activation and attenuates acute inflammatory responses, highlighting its potential as a promising and translationally relevant therapeutic platform for TBI and related neuroinflammatory disorders.

## 1 Introduction

Traumatic brain injury (TBI) is among the leading causes of mortality and disability worldwide, accounting for a substantial proportion of injury-related deaths and long-term neurological impairment (Dawood et al., 2025). Clinical outcomes following TBI are highly heterogeneous and include cognitive deficits, motor dysfunction, and neuropsychiatric complications (Taylor et al., 2017). Globally, TBI affects an estimated ∼70 million individuals annually and involves pathophysiological cascades that extend far beyond the primary insult, contributing to progressive and often lifelong neurological dysfunction (Wu et al., 2021). Importantly, TBI significantly increases the risk of developing chronic neurological disorders, including epilepsy, stroke, and neurodegenerative diseases (Brett et al., 2022).

TBI is a multifaceted pathological event characterized by an initial mechanical injury followed by a cascade of secondary injury processes within the brain parenchyma (Shao et al., 2022). These include disruption of blood-brain barrier (BBB) integrity, excitotoxicity, oxidative stress, endoplasmic reticulum (ER) stress, neuronal dysfunction, axonal degeneration, and persistent neuroinflammation (Shi et al., 2026). The hostile microenvironment created by these processes, characterized by elevated reactive oxygen species, inflammatory mediators, and cellular stress responses, not only exacerbates neuronal injury but also limits the efficacy of therapeutic interventions, including regenerative stem cell transplantation. Therefore, targeting the mechanisms underlying secondary injury remains a critical therapeutic priority.

Neuroinflammation is a central and highly dynamic component of TBI pathology that contributes to both acute damage and chronic neurodegeneration (Holcomb et al., 2025). Microglia, the resident immune cells of the central nervous system (CNS), are key regulators of this response and exhibit remarkable functional plasticity (Loane & Kumar, 2016). In the acute phase, microglia adopt protective roles by clearing cellular debris and secreting neurotrophic and anti-inflammatory factors; however, prolonged activation drives a detrimental pro-inflammatory state characterized by the release of cytokines such as TNF-α, IL-1β, and IL-6, as well as reactive oxygen and nitrogen species (Obukohwo et al., 2024; Simon et al., 2017) (Li et al., 2025). Microglial activation states are broadly categorized into pro-inflammatory M1 and anti-inflammatory/reparative M2 phenotypes, with a well-documented shift toward sustained M1-like activation during chronic phases of injury (Guo et al., 2022; Kumar, Alvarez-Croda, et al., 2016; Wang et al., 2013). In parallel, emerging evidence indicates that glial responses are highly heterogeneous and influenced by developmental stage and cellular context, with distinct astrocyte and microglial subpopulations responding differentially to injury. Thus, therapeutic strategies aimed at reprogramming microglial/macrophage polarization toward a reparative phenotype represent a promising avenue for improving long-term outcomes (Kumar, Barrett, et al., 2016; Wang et al., 2015).

Mitogen-activated protein kinases (MAPKs) are central regulators of cellular stress responses and inflammatory signaling (Moens et al., 2013). This conserved family of serine/threonine kinases, including p38, ERK, and c-Jun N-terminal kinase (JNK) isoforms, is rapidly activated following TBI and governs key processes such as cytokine production, apoptosis, and microglial polarization. Among these, MAPK9 (JNK2) plays a critical role in sustaining pro-inflammatory signaling (Han et al., 2016) and is considered a major regulator of both neuronal death and regeneration caused by neuronal stress (Coffey et al., 2002) and brain injury (Yatsushige et al., 2007). MAPK9 has been identified as a biomarker associated with Alzheimer-related genes (Huang et al., 2025), with potential predictive value for overall survival (Zhang & Kiryu, 2023). Therefore, the modulation of MAPK signaling represents a promising therapeutic approach to limit neuroinflammation and promote tissue repair. Recent advances in gene-editing technologies and nanomedicine have enabled targeted modulation of pathological pathways in the brain. CRISPR/Cas systems provide a powerful approach to selectively disrupt disease-driving genes, while nanoparticle-based delivery platforms offer opportunities to overcome the limitations imposed by the BBB. Intranasal administration has emerged as a non-invasive route for direct nose-to-brain delivery of lipid nanoparticles (LNPs), enhancing CNS targeting while minimizing systemic exposure (Borbolla-Jiménez et al., 2025).

Previously, we demonstrated that LNPs can safely and effectively access the injured brain without exacerbating neuroinflammation (Kara et al., 2025). Building on this, we hypothesized that MAPK9-dependent signaling sustains pro-inflammatory microglial activation after TBI and that CRISPR-mediated MAPK9 editing would lead to sustainable reprogramming of these cells toward a reparative phenotype. To test this, we developed an intranasal LNP-based CRISPR delivery system conjugated to an Iba-1 antibody to selectively target microglia. Using this approach, we show that MAPK9 editing shifts microglial polarization from an M1-like pro-inflammatory state toward an M2-associated reparative phenotype, and reduces pro-inflammatory cytokine production (Quero et al., 2017; Zhang et al., 2019). Collectively, our findings identify targeted genetic modulation of MAPK9 as a promising therapeutic strategy to mitigate neuroinflammation and improve outcomes following TBI.

## 2 Material and Methods

### 2.1 CRISPR sgRNA generation

sgRNA sequences were designed using the Benchling tool to contain a targeting sequence directed to the Mapk9 gene with a 5′-TTTN-3′ PAM compatible with the Cas12a CRISPR enzyme. sgRNA oligonucleotide templates, which included the T7 promoter and the sgRNA reverse complement sequence, were purchased from Eurofins Genomics (Louisville, KY). sgRNA was generated using the MEGAscript™ T7 Transcription Kit (Invitrogen, Carlsbad, CA) according to the manufacturer’s protocol and purified using Oligo Clean & Concentrator™ (Zymo Research, Irvine, CA). The concentration of resulting sgRNA was measured using the RiboGreen® RNA Quantitation Reagent and a Synergy H4 Hybrid Reader (Biotek, Winooski, VT).

### 2.2 CRISPR-LNPs production

LNPs were produced by combining 20 mg S100 soybean phosphatidylcholine (Lipoid, Germany), 8.32 mg cholesterol (Sigma, St. Louis, MO), 3.5 mg DLin-DMA-MC3 (MedChemExpress, Princeton, NJ), 2.24 mg DSPE-PEG(2000) Carboxylic Acid (Avanti Polar Lipids, Alabaster, AL), and a fluorescent dye, 0.2 mg Lissamine™ Rhodamine B 1,2-Dihexadecanoyl-sn-Glycero-3-Phosphoethanolamine (Invitrogen, Waltham, MA) in 4 mL ethanol (Sigma St. Louis, MO). The solvent was placed in a rotary evaporator and evaporated under low pressure (58 mbar) at 100 rpm, in a 41°C water bath to produce a lipid thin film. The resulting thin film was rehydrated with 0.5 mL of sodium acetate buffer (Sigma Aldrich, St. Louis, MO). The CRISPR complex was prepared by mixing 15 μg Cas12a nuclease (Integrated DNA Technologies, Coralville, IA) with 3.75 μg sgRNA in 0.1 mL of sodium acetate buffer and incubating for 15 min at room temperature. The CRISPR complex was then added to 0.125 mL LNP solution, incubated for 15 min at room temperature, and processed using a mini extruder through 0.2 μm filters. CRISPR-LNP size and zeta potential were assessed by dynamic light scattering via a Zetasizer (Malvern, Worcestershire, UK). Encapsulation efficiency was evaluated by separating unbound CRISPR materials from CRISPR-LNPs using an Amicon Ultra 300 kDa centrifugal filter (EMD Millipore, Burlington, MA). The filtrates containing unbound CRISPR complex were measured using the BCA protein concentration assay and confirmed via the Ribogreen® RNA Quantitation Reagent for the sgRNA content. The carboxylic groups on the lipid heads of the resulting CRISPR-LNPs were activated using EDC/NHS according to the manufacturer’s protocol (Thermo Fisher Scientific, Waltham, WA). For Iba-1 attachment, a 3% molar ratio of Iba-1 antibody (Wako, Richmond, VA) was added to the EDC/NHS-activated LNPs to produce Iba-1-conjugated CRISPR-LNPs.

### 2.3 Single-cell RNA-seq data processing

To evaluate the relevance of *Mapk9* in TBI, we analyzed publicly available murine single-cell RNA sequencing (scRNA-seq) data from a CCI injury model obtained from the Gene Expression Omnibus (GEO; accession GSE269748) (Jha et al., 2024). Raw gene expression matrices were processed using Seurat (v5.3.1). Quality control was performed on a per-sample basis using MiQC, retaining cells with a posterior probability ≥ 0.9. Data were normalized using the LogNormalize method and log-transformed, and the top 2,000 highly variable genes were selected for downstream analyses. Batch correction and dataset integration were performed using Harmony (v1.2.3), incorporating sample identity and injury condition as covariates. Principal component analysis (PCA) was conducted on scaled expression values, and the top 41 principal components were used to construct a shared nearest neighbor (SNN) graph. Cells were clustered using the Louvain algorithm (resolution = 0.5), yielding 25 transcriptionally distinct clusters. Low-dimensional visualization was performed using Uniform Manifold Approximation and Projection (UMAP) based on Harmony-corrected embeddings. Cell-type annotation was performed using SingleR, followed by manual refinement based on canonical marker genes and reference datasets. Differential gene expression analysis was conducted using the Wilcoxon rank-sum test implemented in Seurat, with acceleration via the presto R package. Genes with an adjusted *P* value < 0.05 (Benjamini–Hochberg correction) were considered significant and ranked by log fold change. Gene expression patterns were visualized using UMAP feature plots derived from log-normalized data. The proportion of *Mapk9*-expressing cells (nPos) was calculated for each annotated cell population under naïve and CCI conditions, and cell-type composition was quantified as the percentage of total cells per condition.

### 2.4 CRISPR-LNPs assessment *in vitro*

Primary macrophages were isolated from mouse bone marrow and cultured in macrophage differentiation media for 7 days to obtain naïve macrophages. Undifferentiated primary macrophages were seeded on 8/16-well chamber slides (Nunc™ Lab-Tek™) with a 30,000 cells/cm^2^ seeding density. To assess macrophage differentiation, 2 μL of CRISPR-LNPs was added to each well of primary macrophages seeded in 8-well chamber culture slides and incubated for 24 h. After incubation, the culture medium was changed, and cells were stimulated with 50 ng/mL IL-4 and 50 ng/mL M-CSF to induce *M2* or 50 ng/mL LPS and 25 ng/mL IFN-γ to induce *M1* differentiation. After 48 h of incubation, cells were fixed with 4% paraformaldehyde for 30 min at 4°C and stained with 2.5 μg/mL rat anti-mouse CD80 primary antibody (Thermo Fisher Scientific, Waltham, MA), FITC-rat anti-mouse CD206 antibody (Abcam, Cambridge, UK), and Alexa Fluor 568-goat anti-rat IgG secondary antibody for macrophage phenotype analysis. Cells were analyzed using a Nikon A1 confocal microscope (Nikon Inc., Melville, NY, USA), and fluorescence intensity was assessed with NIS-Elements software (Nikon Inc.).

### 2.5 RNA extraction and qPCR

Total RNA was extracted from macrophages using TRIzol™ Reagent (Invitrogen, cat. #15596026) according to the manufacturer’s instructions. One microgram of each sample’s isolated RNA served as a template for reverse transcription into complementary DNA (cDNA), performed with the iScript cDNA Synthesis Kit (Bio-Rad, cat. #1708891) under the following conditions: 25°C for 5 min, 46°C for 20 min, and 95°C for 1 min. The cDNAs were amplified with SsoAdvanced Universal SYBR Green Supermix using the CFX384 Touch Real-Time PCR Detection System (Bio-Rad). The PCR protocol was set up as follows: an initial denaturation step at 95°C for 3 min, followed by 40 cycles of denaturation at 95°C for 10 s, annealing at 50°C for 30 s, and extension at 72°C for 30 s. Gene expression levels were normalized to the housekeeping gene, β-actin. Relative differences in expression were measured using the comparative threshold cycle (2^-ΔΔCt^) method.

### 2.6 Mice and Traumatic brain injury model

Adult male (12-week-old) C57BL/6J mice (Jackson Laboratories, Bar Harbor, ME) were housed at the Houston Methodist Research Institute (HMRI) animal facility. Mice were maintained on a 12 h light-dark cycle and had *ad libitum* access to food and water. All experiments were approved by the Institutional Animal Care and Use Committee (IACUC) at HMRI. Studies were conducted in accordance with the NRC Guide for the Care and Use of Laboratory Animals. Mice were anesthetized with isoflurane (3% for induction, 1.5-2% for maintenance) before and during surgery. Controlled cortical impact (CCI) injury was induced on the left hemisphere, targeting the primary motor and somatosensory cortex using an electromagnetic Impact One stereotaxic impactor (Leica Biosystems, Buffalo Grove, IL, USA). The impact site was localized 2 mm lateral and 2 mm posterior to Bregma, using a 3-mm-diameter flat impact tip, with an impact velocity of 3.25 m/s, and an impact depth of 1.5 mm. Sham mice underwent all procedures, including anesthesia, but did not undergo CCI.

### 2.7 Microglial targeting of CRISPR-LNPs

We next evaluated *in vivo* biodistribution of fluorescently labeled Iba-1-CRISPR-LNPs following retro-orbital (100 μL) or intranasal (25 μL per nostril) administration under anesthesia 30 min post-injury, at a dose of 20 mg/kg CRISPR. After 24 h of TBI, mice were euthanized, and their organs, including the brain, heart, lungs, spleen, liver, kidneys, and blood, were collected for further analysis.

### 2.8 Immunofluorescence analysis

Brains were fixed in 4% paraformaldehyde overnight. After 24 h of fixation, they were cryopreserved in a 30% sucrose solution for 48 h. Serial free-floating coronal brain sections, 16 μm thick, were prepared at the level of the dorsal hippocampus for immunohistochemical analysis. Sections were processed using immunohistochemistry protocols that included three consecutive 5-min washes in PBS-T (PBS containing 0.5% Triton X-100). Sections were incubated with 5% normal goat serum (NGS) in PBS-T for 1 h at room temperature to block nonspecific binding. Brain sections were then incubated overnight at 4°C in a solution of 3% NGS in PBS-T, containing primary antibodies: anti-rabbit Iba-1 (Wako, cat. #019-19741) at 1:500 to label microglia and macrophages, anti-mouse NeuN (Millipore Sigma, cat. #MAB377) at a 1:200 dilution to detect neurons. After incubation, tissue sections were washed three times with PBS-T for 5 min and then incubated for 2 h at room temperature with the corresponding secondary antibodies: Goat anti-Rabbit Alexa Fluor™ 488 (Invitrogen, cat. #A11034) and Goat anti-Mouse Alexa Fluor™ 488 (Invitrogen, cat. #A21121), at a 1:1000 dilution. The sections were then rinsed three times for 5 min each in PBS and incubated in PBS containing DAPI (1:50,000; Sigma-Aldrich, St. Louis, MO) for nuclear counterstaining. The sections were rinsed with distilled water and mounted using Fluoro-Gel with Tris buffer.

### 2.9 Fluorescent in situ hybridization with immunohistochemical labeling

Coronal brain sections were mounted on gelatin-coated glass slides (Superfrost Plus, Fisher Scientific, cat. #12-550-15) and kept at -80°C until use. Fluorescent in situ hybridization (FISH) was performed using the RNAscope™ 2.5 HD Reagent Kit-RED, following the manufacturer’s instructions (Advanced Cell Diagnostics, cat. #322350), as previously described (Flinn et al., 2026; Villapol et al., 2017). Brain tissue sections were dehydrated through a graded ethanol series (50%, 70%, and 100%; twice for 5 min each). The sections were then incubated in Pretreatment 1 solution for 10 min, followed by boiling in Pretreatment 2 solution for 2 min. Finally, the slides were incubated in Pretreatment 3 solution for 30 min prior to hybridization. Sections were incubated at 40°C for 2 h for hybridization with the specific target probes separately: *mus musculus* MAPK9 (Advanced Cell Diagnostics, cat. #486031), *mus musculus* IL-1β (Advanced Cell Diagnostics, cat. #316891), and *mus musculus* TNF-α (Advanced Cell Diagnostics, cat. #311081). Additionally, the negative control probe (Advanced Cell Diagnostics, cat. #310043) and the positive control probe (Advanced Cell Diagnostics, cat. #313911) were applied and incubated for 2 h at 40°C. The amplification steps were performed based on the manufacturer’s instructions.

### 2.10 Cell death assay and lesion volume measurements

To assess cell death, brain sections were analyzed for DNA strand breaks using Terminal deoxynucleotidyl transferase dUTP nick end labeling (TUNEL) from the Fluorescence In Situ Cell Death Detection kit (Roche Diagnostics, Indianapolis, IN) according to the manufacturer’s protocol. Brain sections were stained with cresyl violet for lesion volume analysis. To visualize injury-related regions, an average of 10-12 coronal brain sections, evenly spaced from 0 to - 2.70 mm relative to bregma, were chosen for cresyl violet staining. Sections were mounted on gelatin-coated glass slides (SuperFrost Plus, Thermo Fisher Scientific, IL) and immersed in a freshly prepared 0.5% cresyl violet solution (Sigma-Aldrich, St. Louis, MO) prepared in distilled water and filtered before use. Next, the slides were dehydrated through graded ethanol solutions (100%, 95%, 70%, and 50%) for 2 min each, then cleared in xylene twice for 2 min each. The sections were then sealed with Permount mounting medium (Thermo Fisher Scientific) to ensure preservation. The lesion volume was calculated as a percentage of the lesion area. The data from each slide, comprising 10-12 brain sections, were then averaged in ImageJ.

### 2.11 Quantitative analysis of immunolabeled images

All histological images were obtained using a Slideview VS200 Universal Whole Slide Imaging Scanner (Evident, USA) and a confocal imaging system (Leica Microsystems, Deerfield, IL, USA). We used unbiased, standardized sampling techniques to measure immunoreactivity in cortical tissue areas showing positive staining. For proportional area measurements, microglia/macrophage Iba-1 immunoreactivity was expressed as the proportion of the target region covered by immunohistochemically stained cellular profiles. To quantify Iba-1+ and TUNEL+ cells in the injured cortex, we analyzed an average of four coronal sections from the lesion epicenter (-1.34 to -2.30 mm from Bregma) for each mouse.

### 2.12 Rotarod test

The rotarod test is a method for assessing sensorimotor function in mice. For this evaluation, we used the Rotamex 5 system from Columbus Instruments (Columbus, OH) and followed the protocol as previously described (Baudo et al., 2024; Zinger et al., 2021). Each mouse underwent a 2-day training regimen consisting of 3 daily trials, during which they were allowed to explore the testing apparatus for a few min before testing began. Then, the rod began to rotate, with the drum’s speed gradually increasing from 4 to 40 rpm. Each trial ended when the mouse fell off the rotarod or after 5 min, and the latency to fall was recorded. The mean value of these measurements served as the baseline assessment for each mouse. 1-day post-injury (dpi), mice were also tested on the rotarod.

### 2.13 AST activity assay

AST activity in serum and liver was measured using the Aspartate Aminotransferase Activity Assay Kit (Abcam, cat. #ab105135) following the manufacturer’s instructions. Blood samples from each mouse were collected, and serum was obtained through centrifugation at 4000 rpm for 20 min at 4°C. 50 mg of liver tissue from each mouse was homogenized in assay buffer, and the supernatant was collected after centrifuging at 13500 rpm for 15 min 4°C. Serum and liver samples were diluted 1:10 in assay buffer and added to the wells of a 96-well plate along with the colorimetric assay reaction mix. The plate was incubated for 60 min at 37°C, and the absorbance was measured at 450 nm. AST activity for each sample was calculated using the standard curve based on glutamate standard readings.

### 2.14 Western Blot

Serum samples (1:5 v/v) were mixed with 4x Laemmli sample buffer (Bio-Rad, cat. #1610747) and then incubated at 100°C for 5 min. The proteins were separated by SDS-PAGE with a 4%-20% gradient and subsequently transferred to polyvinylidene difluoride (PVDF) membranes (Bio-Rad, cat. #1620177). The membranes were blocked with a 5% w/v skim milk powder solution in PBS-Tween 20 buffer for 1 h at room temperature. Serum amyloid A1/A2 (SAA) expression was detected using a specific SAA antibody (1:1000 dilution, R&D Systems, cat. #AF2948) and an HRP-conjugated secondary antibody. Immunoblots were visualized with Clarity Western ECL (Bio-Rad, cat. #1705061) on a ChemiDoc MP imaging system (Bio-Rad), and bands were quantified using a densitometer in ImageJ.

### 2.15 Organs paraffin embedding and Hematoxylin and Eosin staining

Kidney, spleen, lung, heart, and liver were sampled, fixed in 4% paraformaldehyde for 2 days, and transferred to 70% ethanol. Tissues were processed using the manufacturer’s standard protocols in a Shandon Exelsion ES Tissue Processor and embedded in paraffin on a Shandon HistoCenter Embedding System. Following dehydration with 95% ethanol twice for 30 min, samples were soaked in xylene for 1 h at 60-70°C, and then in paraffin for 12 h. 0.5 mL of 95% ethanol was used to dehydrate the mouse organs. Tissues were stained with hematoxylin solution for 6 h at 60-70°C, then rinsed with water until the water was colorless. Next, the tissues were placed in 10% acetic acid and 85% ethanol twice for 2 min each, then rinsed with tap water. Nuclei were blue-fluorescently stained using NucBlue Fixed Cell read probes (Thermo Fisher Scientific, IL, USA) for 10 min, then rinsed again with water. Hematoxylin and Eosin staining was performed, and slides were stored at room temperature. Representative images of all organs were taken on the microscope at 20x.

### 2.16 Statistical analysis

A one-way analysis of variance (ANOVA) followed by Tukey’s multiple-comparison test was used for microglia-targeting studies, rotarod test, AST activity assay, and Western Blot analysis. Immunofluorescence staining, including Iba-1+ and TUNEL+ cells, was analyzed using one-way ANOVA. Student’s t-test was used for the qPCR assay, RNAscope data, and lesion volume analysis. All mice were randomly assigned to experimental groups, and the researchers remained blinded to the treatments throughout the entire study. Data are shown as the mean ± SEM. Statistical analysis was performed with GraphPad Prism 10 (GraphPad Software, San Diego, CA, USA) for multiple groups, assuming a normal data distribution. Significance levels are indicated as *p < 0.05, **p < 0.01, ***p < 0.001, and ****p < 0.0001.

## 3 Results

### 3.1 Engineering and characterization of Iba-1-targeted CRISPR-LNPs

We engineered LNPs encapsulating CRISPR-Cas12a/Cpf1 targeting M2-polarization genes and functionalized their surface with an Iba-1 antibody using EDC/NHS chemistry to enhance microglial targeting (**Fig. 1A**). Rhodamine labeling enabled tracking of nanoparticle biodistribution. Physicochemical characterization revealed that both CRISPR-LNPs and Iba-1-CRISPR-LNPs exhibited a stable negative zeta potential, with a modest shift following antibody conjugation (**Fig. 1B**), consistent with reduced nonspecific protein adsorption and favorable circulation properties. Dynamic light scattering showed a uniform size distribution (∼160 nm) with no significant change after Iba-1 modification (**Fig. 1C**). The CRISPR system is designed to shift microglial activation from a proinflammatory M1 phenotype toward an anti-inflammatory M2 state (**Fig. 1D**). To validate these effects *in vivo*, the experimental design employs a TBI animal model, with motor behavior and outcome measures assessed at 1 dpi (**Fig. 1E**).

**Figure 1.**
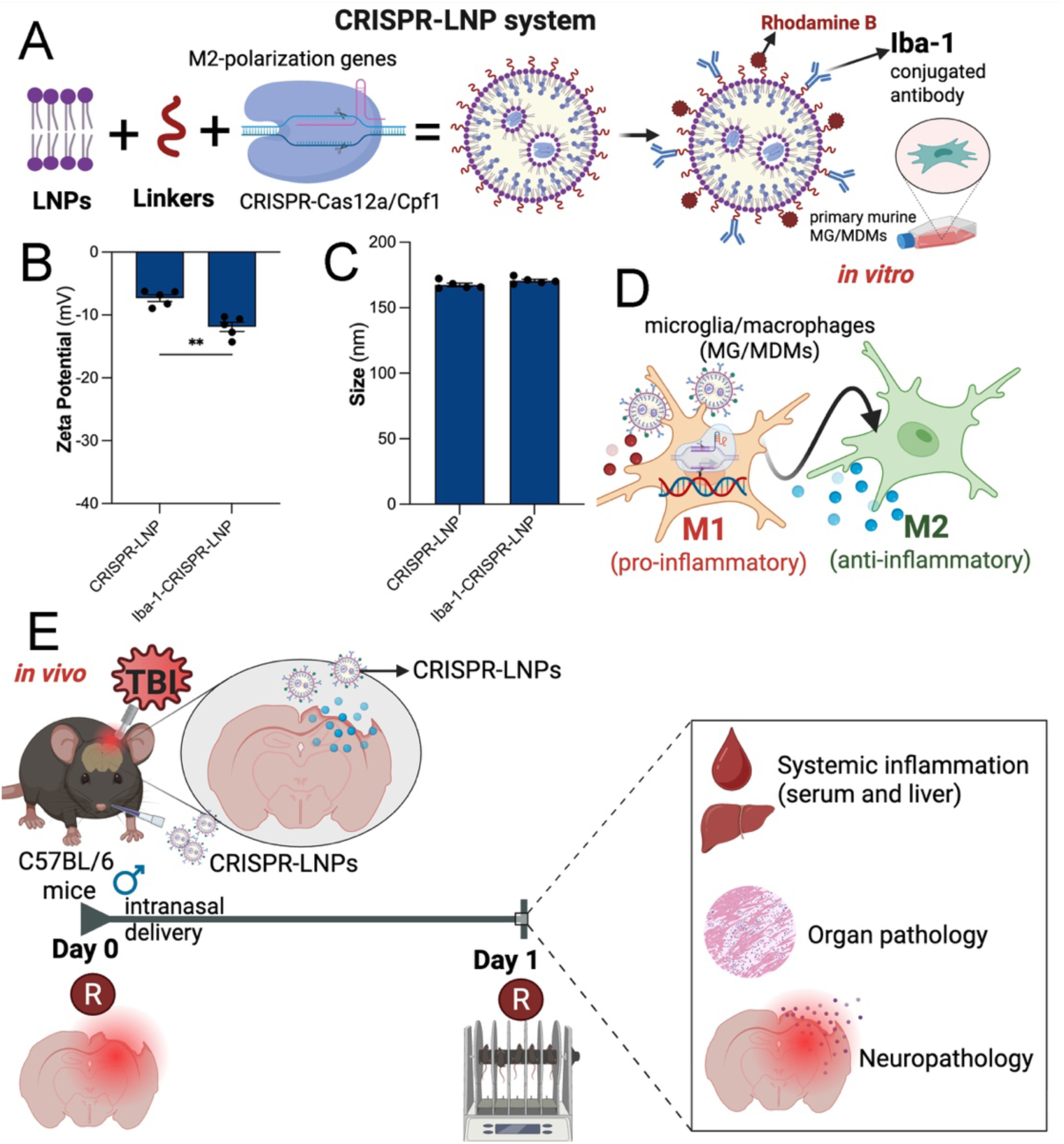
CRISPR-LNP platform for microglia/macrophage reprogramming and *in vivo* evaluation after TBI. (**A**) Schematic of CRISPR-loaded lipid nanoparticles (CRISPR-LNPs). Lipid nanoparticles (LNPs) are assembled with linker chemistry and CRISPR-Cas12a/Cpf1 components targeting M2-polarization genes. Surface functionalization with Rhodamine B and Iba-1-conjugated antibodies enables targeting of microglia/macrophages (MG/MDMs) and *in vitro* validation in primary murine cells. (**B**) Zeta potential measurements of CRISPR-LNPs and Iba-1-CRISPR-LNPs, showing surface charge modification following antibody conjugation. (**C**) Particle size distribution of CRISPR-LNPs and Iba-1-CRISPR-LNPs, demonstrating maintained nanoscale size after functionalization. (**D**) Mechanism of action: CRISPR-LNP uptake by MG/MDMs induces gene editing to shift polarization from a pro-inflammatory M1 phenotype to an anti-inflammatory M2 phenotype. (**E**) *In vivo* experimental design. TBI is induced in male C57BL/6 mice followed by intranasal administration of CRISPR-LNPs. Downstream analyses include systemic inflammation (serum and liver), organ pathology, and brain neuropathology at 1-day post-TBI.

### 3.2 CRISPR-LNPs promote LPS-induced primary macrophage transformation from M1-like to M2-like phenotype

We treated murine primary macrophages with Iba-1-CRISPR-LNPs for 48 h. We observed that CRISPR-LNPs significantly drove expression of M2-associated marker genes (CD206+) upon MAPK9 targeting, including NTN1 and SOCS3, even under strongly pro-inflammatory conditions (LPS+IFNγ), which normally drive M1-like polarization (**Fig. 2A**). Over 70% of untreated control macrophages expressed CD80 and less than 25% expressed CD206, while nearly 100% of the cells treated with MAPK9 or SOCS3 CRISPR-LNPs expressed CD206. CRISPR-LNP treatment prevented M1-associated profile and stabilized an M2-like, pro-repair phenotype (**Fig. 2B**). Treatment of macrophages with CRISPR-LNPs targeting MAPK9 reduced mRNA levels of pro-inflammatory cytokine genes (MCP-1, MIP-1α, MIP-1β, CCL5, CXCL1, IL-1β) significantly, demonstrating that our system reprogrammed microglia, shifting the pro-inflammatory M1 phenotype to an anti-inflammatory M2 phenotype *in vitro* (**Fig. 2C**).

**Figure 2.**
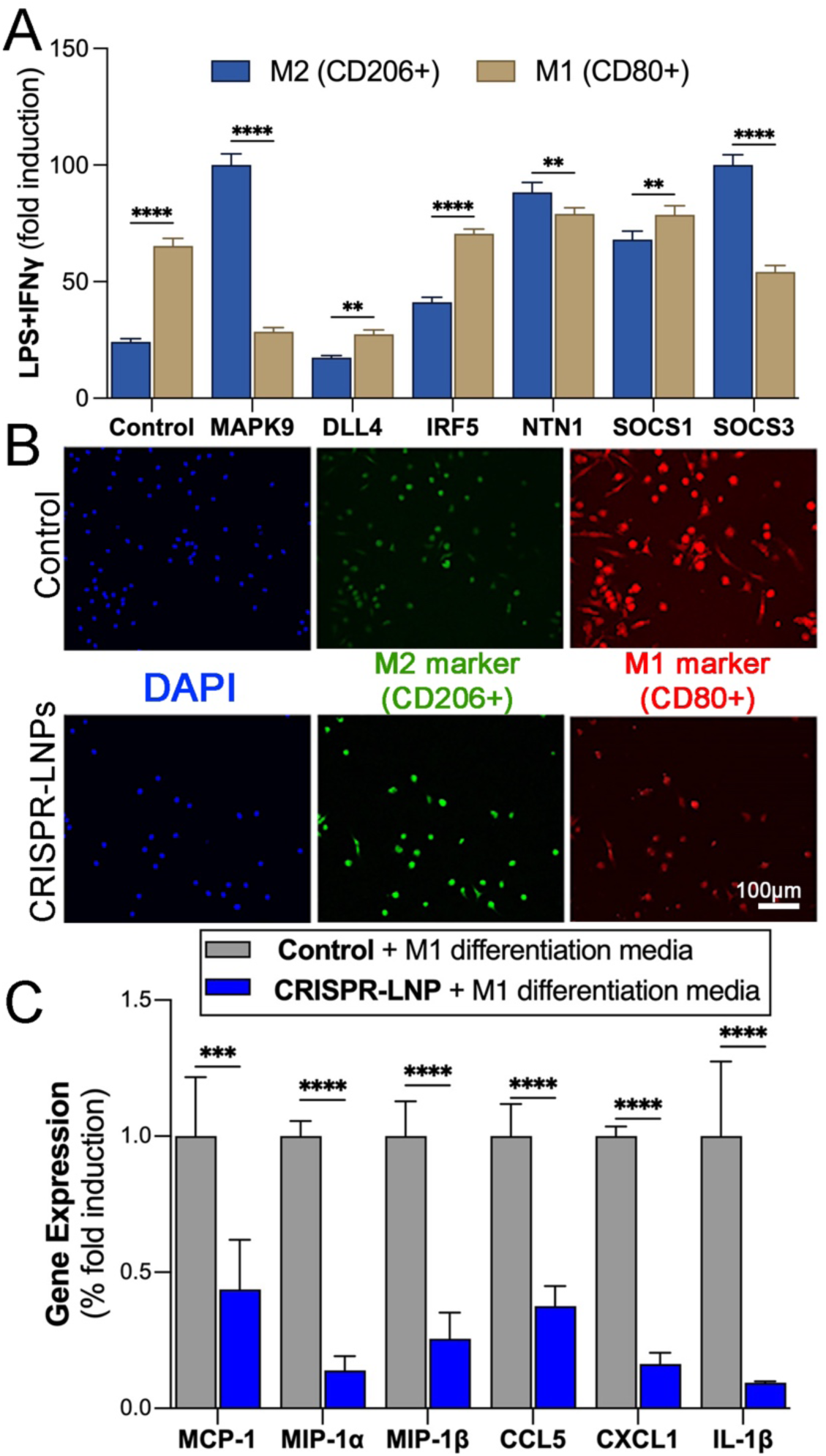
CRISPR-LNPs reprogram microglia/macrophage polarization and suppress pro-inflammatory responses *in vitro*. (**A**) Gene modulation associated with macrophage polarization following CRISPR-LNP treatment under pro-inflammatory conditions (LPS+IFNγ). Percentage of cells expressing M2-associated marker (CD206+) and M1-associated marker (CD80+) genes is shown after CRISPR treatment targeting including *MAPK9, DLL4, IRF5, NTN1, SOCS1,* or *SOCS3* genes. CRISPR-LNP treatment promotes M2-associated profiles while reducing M1-associated responses for selected targets. (**B**) Representative immunofluorescence images of microglia/macrophages under control conditions or treated with CRISPR-LNPs in the presence of M1 differentiation media. Nuclei are labeled with DAPI (blue), M2 marker CD206 (green), and M1 marker CD80 (red). CRISPR-LNP treatment increases CD206+ cells and reduces CD80+ cells compared to control. Scale bar: 100 μm. (**C**) Quantification of pro-inflammatory gene expression (MCP-1, MIP-1α, MIP-1β, CCL5, CXCL1, and IL-1β) in control versus CRISPR-LNPs-treated cells under M1 differentiation conditions. CRISPR-LNPs targeting MAPK9 significantly reduce the expression of inflammatory mediators compared to control. Data are presented as mean ± SEM. **p < 0.01, ***p < 0.001, ****p < 0.0001.

### 3.3 Single-cell RNA-seq analysis from Gene Expression Omnibus (GEO) database

To ensure consistency with our experimental framework, only male samples collected at 24 h post-CCI were included in the analysis (**Fig. 3**). Major cell populations identified included microglia, bone marrow-derived monocytes/macrophages (BMD Mono/Macs), neutrophils, dendritic cells, T/NK cells, B cells, astrocytes, oligodendrocytes, neurons, ependymal cells, endothelial cells, and pericytes. Single-cell RNA-seq analysis identified distinct cellular populations across naïve and CCI conditions at 1 dpi, including microglia, BMD Mono/Macs, neutrophils, dendritic cells, and multiple neural cell types (**Fig. 3A**). UMAP clustering revealed clear segregation of immune and resident brain cell populations, with microglia forming the dominant cluster and peripheral immune populations (e.g., neutrophils and BMD Mono/Macs) occupying distinct transcriptional spaces. Cellular composition analysis demonstrated a marked shift following CCI (**Fig. 3B**). While microglia remained the predominant population in both conditions, there was a notable increase in infiltrating BMD Mono/Macs and neutrophils after injury, consistent with an acute inflammatory response. Other cell types showed relatively minor proportional changes at this early time point. Expression of Mapk9 was enriched primarily within myeloid populations and displayed condition-dependent differences. In microglia (**Fig. 3C**), the proportion of Mapk9-positive cells increased from 4.39% in naïve to 8.01% in CCI, indicating an injury-associated upregulation within resident immune cells. A similar trend was observed in BMD Mono/Macs (**Fig. 3E**), where Mapk9-positive cells increased in both number and proportion following CCI (4.9% to 6.93%), suggesting activation of infiltrating myeloid populations. In contrast, astrocytes exhibited a reduction in Mapk9-positive cells after injury (5.59% to 1.79%; **Fig. 3D**), indicating that Mapk9 expression is not broadly induced across all glial populations and may be more specific to myeloid lineage responses. Oligodendrocytes showed a modest decrease in Mapk9-positive cells (11.35% to 7.43%; **Fig. 3F**), although expression remained detectable. Notably, Mapk9 expression was absent in naïve neutrophils but emerged following CCI (0% to 0.93%; **Fig. 3G**), consistent with recruitment of activated peripheral immune cells expressing this kinase. Neurons displayed relatively stable Mapk9 expression between conditions (7.09% vs. 7.96%; **Fig. 3H**), suggesting that neuronal Mapk9 levels are not strongly altered at this early post-injury stage. Overall, these data indicate that Mapk9 expression is preferentially associated with myeloid cell activation following TBI, particularly in microglia and infiltrating macrophages, with limited or variable changes observed in non-immune brain cell populations.

**Figure 3.**
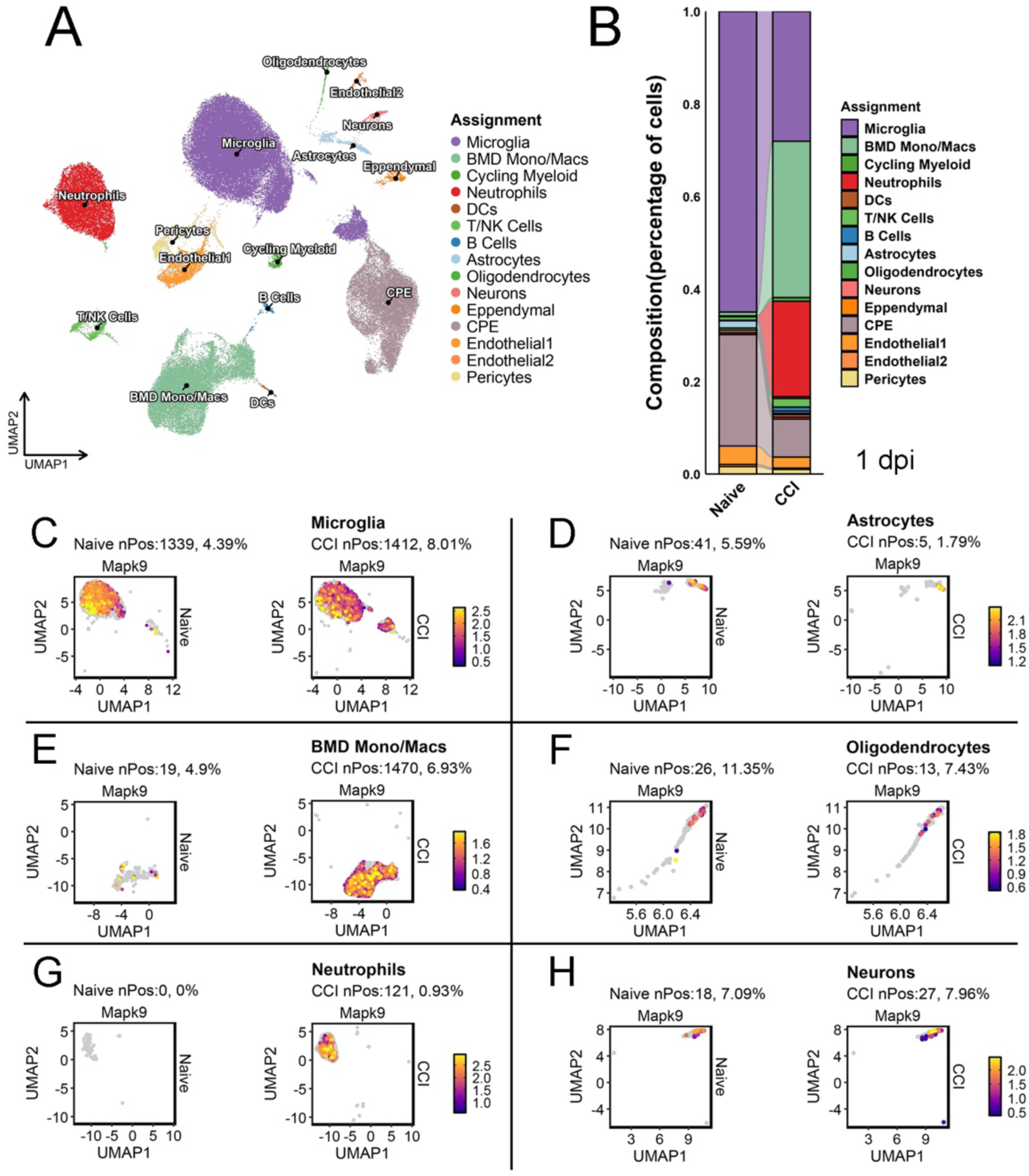
Single-cell RNA-seq reveals cell-type–specific Mapk9 expression following TBI. **(A)** UMAP visualization of integrated single-cell RNA-seq data from naïve and controlled cortical impact (CCI) mouse brains at 1 day post-injury (1 dpi) identifies major cell populations, including microglia, bone marrow-derived monocytes/macrophages (BMD Mono/Macs), neutrophils, dendritic cells (DCs), T/NK cells, B cells, astrocytes, oligodendrocytes, neurons, ependymal cells, choroid plexus epithelial (CPE) cells, endothelial cells, and pericytes. (**B**) Stacked bar plots show relative cellular composition, indicating an increased representation of myeloid populations following CCI. (**C-H**) Feature plots of Mapk9 expression across selected cell types demonstrate cell-type-specific and injury-dependent changes, with an increased proportion of Mapk9-positive cells in microglia (4.39% naïve vs. 8.01% CCI) and BMD Mono/Macs (4.9% vs. 6.93%), emergence of Mapk9 expression in neutrophils after injury (0% vs. 0.93%), and relatively stable expression in neurons (7.09% vs. 7.96%). In contrast, astrocytes (5.59% vs. 1.79%) and oligodendrocytes (11.35% vs. 7.43%) show reduced proportions of Mapk9-positive cells following CCI. Percentages indicate the fraction of Mapk9-positive cells (nPos) within each cell type for each condition.

### 3.4 Intranasal delivery enhances microglial targeting and MAPK9 editing *in vivo*

We next evaluated the *in vivo* biodistribution and cellular targeting of fluorescently labeled Iba-1-CRISPR-LNPs following retro-orbital or intranasal administration. Confocal imaging of cortical sections revealed CRISPR-LNP signal (red) in both delivery conditions, with visible association to Iba-1+ microglia/macrophages (green). Qualitatively, intranasal delivery showed more frequent and spatially concentrated co-localization with Iba-1+ cells compared to retro-orbital administration (**Fig. 4A**). Quantitative analysis confirmed this observation, demonstrating that intranasal delivery significantly increased the proportion of CRISPR-LNP+ signal within Iba-1+ cells, while reducing association with Iba-1- populations (**Fig. 4B**), indicating improved targeting specificity. High-resolution cortical imaging further supported these findings. In Iba-1-labeled sections (**Fig. 4C-E**), the CRISPR-LNP signal was broadly distributed and frequently overlapped with microglial cells. Merged images **(Fig. 4E**) showed clear but partial co-localization, consistent with preferential uptake by microglia/macrophages rather than exclusive targeting. In parallel, neuronal labeling (**Fig. 4F-H**) demonstrated that the CRISPR-LNP signal was also detectable in NeuN+ cells. However, merged images (**Fig. 4H**) indicated comparatively lower colocalization with neurons than with microglia, suggesting that neuronal uptake occurs but is not the dominant targeting outcome. Overall, these data indicate that intranasal administration enables broad cortical penetration of CRISPR-LNPs, with enriched targeting to microglial cells alongside measurable, but lower, neuronal uptake. Importantly, RNAscope analysis at 1 dpi showed that intranasal CRISPR-LNP treatment markedly reduced cortical MAPK9 mRNA levels in CCI mice compared to vehicle-treated controls, confirming successful *in vivo* gene modulation (**Fig. 4I-J**, **i1, j1**).

**Figure 4.**
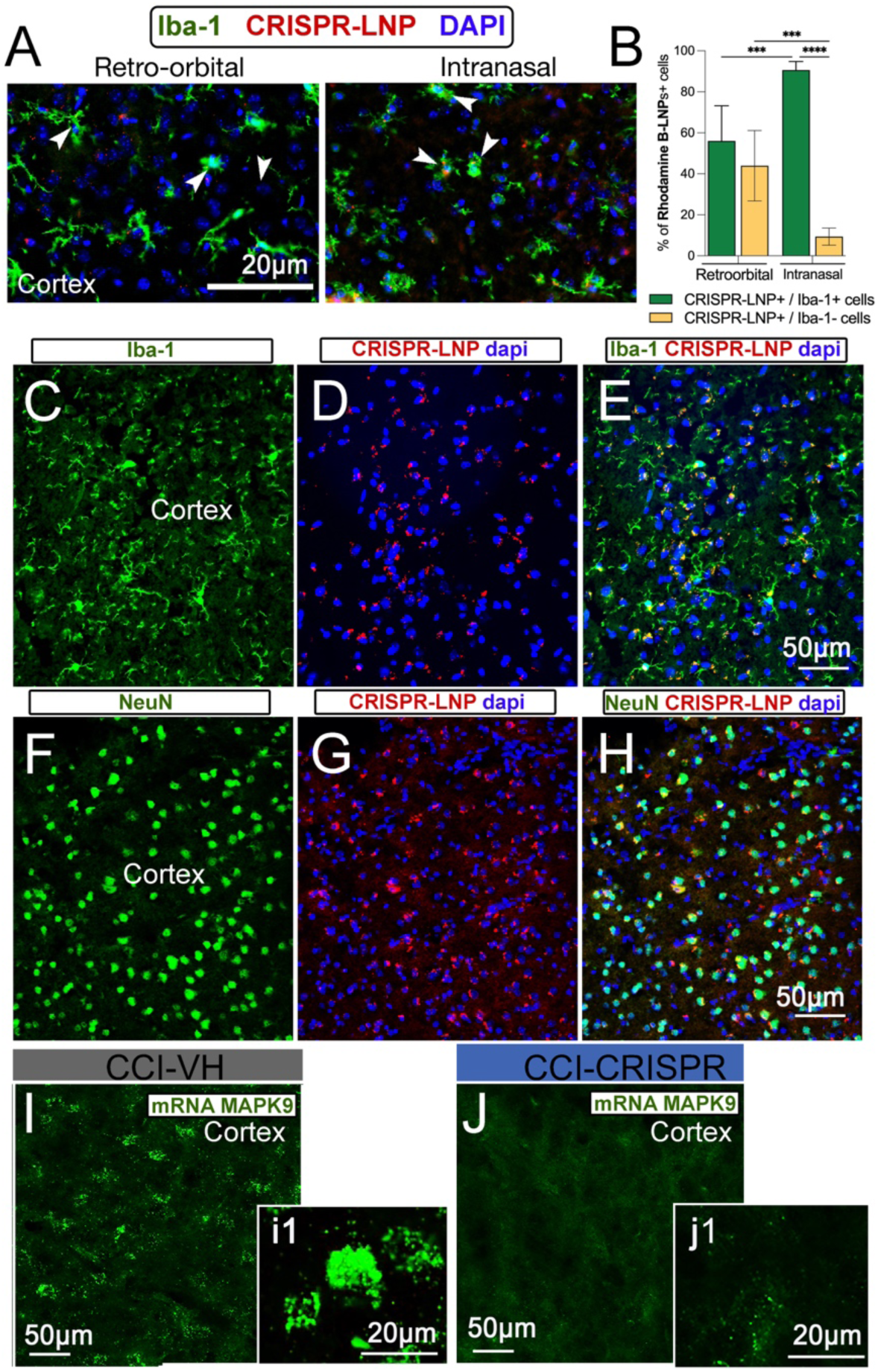
Intranasal delivery enhances brain targeting of CRISPR-LNPs and reduces MAPK9 expression after TBI. (**A**) Representative confocal images of cortical sections showing uptake of Rhodamine B-labeled CRISPR-LNPs (red) in Iba-1+ microglia (green) with DAPI nuclear staining (blue) following retro-orbital (left, 100 μL) or intranasal (right, 25 μL) administration. Arrowheads indicate co-localization of CRISPR-LNPs within microglia. Scale bar: 20 µm. (**B**) Quantification of CRISPR-LNPs+ cells in the cortex, showing the percentage of uptake in Iba-1+ versus Iba-1- cells after retro-orbital and intranasal delivery. Intranasal administration significantly increases microglial targeting. Data are mean ± SEM; ***p < 0.001, ****p < 0.0001. (**C-E**) Representative images of cortical microglia labeled with Iba-1 (**C**), CRISPR-LNPs with DAPI (**D**), and merged image (**E**), demonstrating robust uptake of CRISPR-LNPs in Iba-1+ cells. Scale bar: 50 µm. (**F-H**) Representative images of neurons labeled with NeuN (**F**), CRISPR-LNPs with DAPI (**G**), and merged image (**H**), showing comparatively lower neuronal uptake of CRISPR-LNPs. Scale bar: 50 µm. (**I-J**) RNAscope analysis of MAPK9 mRNA expression in the cortex. Insets (**i1, j1**) show high-magnification views of MAPK9 transcripts (green puncta). Compared to vehicle-treated controls (CCI-VH, **I**), intranasal CRISPR-LNP treatment (CCI-CRISPR, **J**) markedly reduces MAPK9 mRNA levels. Scale bars: 20 µm (insets), 50 µm.

### 3.5 CRISPR-LNPs attenuate microglial activation after TBI

To assess therapeutic efficacy, we examined microglial activation in the peri-contusional cortex at 1 dpi. Iba-1 immunostaining revealed a significant reduction in total Iba-1+ cells in CRISPR-treated CCI mice compared to vehicle (Vh)-treated CCI mice (**Fig. 5A-B**). Morphological analysis further showed that CRISPR-LNP treatment selectively reduced hypertrophic activated microglial phenotypes (**Fig. 5D**), without altering the population of ramified (resting) microglia (**Fig. 5C**) or amoeboid (phagocytic) microglia (**Fig. 5E**). These findings indicate a shift away from the pro-inflammatory M1-like state. Lesion volume was not significantly altered by treatment (**Fig. 5F-G**), suggesting that early effects are driven primarily by immunomodulation rather than structural changes. RNAscope analysis exhibited a non-significant reduction in pro-inflammatory cytokine expression, but a trend, including TNF-α and IL-1β, in CRISPR-treated mice compared with controls (**Fig. 5H-M**). Additionally, TUNEL staining revealed a trend toward reduced cell death, although this did not reach statistical significance at this early time point (**Fig. 5O-Q**). Motor performance assessed by rotarod showed no significant differences between CRISPR- and Vh-treated CCI mice at 1 dpi (**Fig. 5N**). As expected, both groups exhibited impaired performance relative to baseline, reflecting acute injury effects.

**Figure 5.**
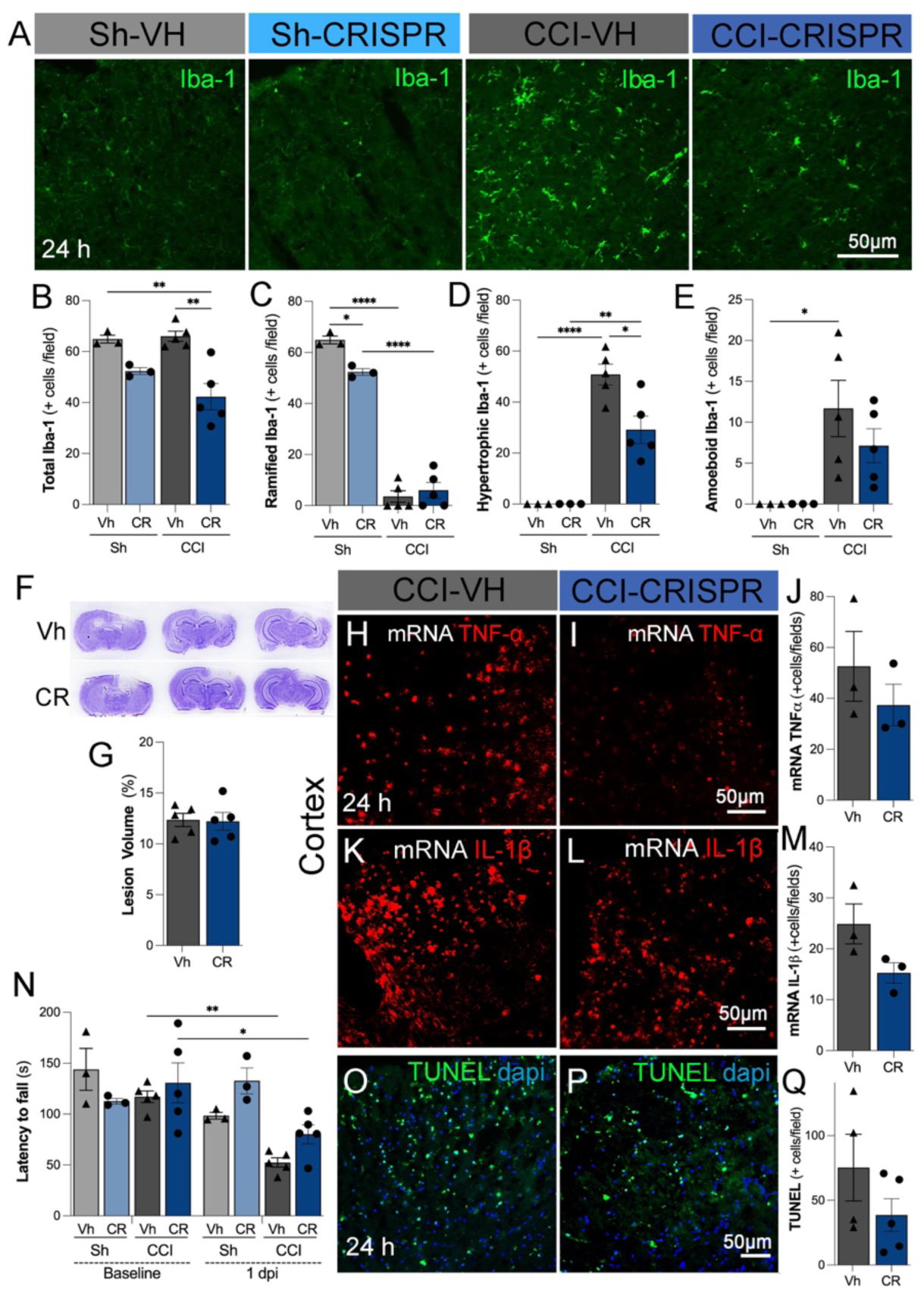
CRISPR-LNP treatment attenuates microglial activation, neuroinflammation, and cell death following TBI. (**A**) Representative immunofluorescence images of Iba-1+ microglia/macrophages in sham (Sh) and controlled cortical impact (CCI) mice treated with vehicle (VH) or CRISPR-LNPs (CRISPR). CCI induces marked microglial activation, which is reduced by CRISPR-LNP treatment in the somatosensory cortex. (**B-E**) Quantification of microglial populations per field. (**B**) Total Iba-1+ cells, (**C**) ramified (resting) microglia, (**D**) hypertrophic (activated) microglia, and (**E**) amoeboid (phagocytic) microglia. CRISPR-LNP treatment reduces total and activated microglial phenotypes following CCI while partially preserving homeostatic morphology. (**F**) Representative cresyl violet-stained brain sections showing lesion morphology in VH- and CRISPR-treated animals after CCI. (**G**) Quantification of lesion volume (%), showing no significant difference between VH- and CRISPR-LNPs-treated groups. (**H-I**) Representative images of *TNF-α* mRNA expression in the cortex of CCI mice treated with VH or CRISPR-LNPs. (**J**) Quantification of *TNF-α* mRNA+ cells per field, showing reduced inflammatory gene expression following CRISPR-LNP treatment. (**K-L**) Representative images of *IL-1β* mRNA expression (red) in cortex sections from CCI-VH and CCI-CRISPR groups. (**M**) Quantification of *IL-1β* mRNA+ cells per field, demonstrating a reduction with CRISPR-LNP treatment. (**N**) Behavioral assessment (latency to fall, rotarod test) at baseline and 1-day post-injury (1 dpi). CRISPR-LNPs-treated mice show improved motor performance compared to VH-treated CCI animals. (**O-P**) Representative TUNEL staining (green) with DAPI (blue) in cortex sections, indicating apoptotic cell death in CCI-VH and reduced cell death in CCI-CRISPR groups. (**Q**) Quantification of TUNEL+ cells per field, showing decreased apoptosis following CRISPR-LNP treatment. Scale bar: 50 µm. Data are presented as mean ± SEM. *p < 0.05, **p < 0.01, ****p < 0.0001.

### 3.6 CRISPR-LNPs reduce systemic inflammation without inducing toxicity

To evaluate systemic effects, we measured SAA, an acute-phase reactive protein that increases following TBI. SAA levels were significantly elevated in CCI mice and were markedly reduced by CRISPR-LNP treatment compared to vehicle controls (**Fig. 6A-B**), indicating attenuation of peripheral inflammation. Serum AST levels were elevated in CCI mice relative to sham controls but were not significantly different between CRISPR- and Vh-treated groups (**Fig. 6C**). Liver AST activity remained unchanged across all groups (**Fig. 6D**). Histopathological analysis of the spleen, heart, kidney, lung, and liver revealed no treatment-related abnormalities in CRISPR-LNP-treated animals compared to controls (**Fig. 6E**). No differences in body weight or general behavior were observed. Overall, these results show that CRISPR-LNP therapy has a safe systemic profile *in vivo*.

**Figure 6.**
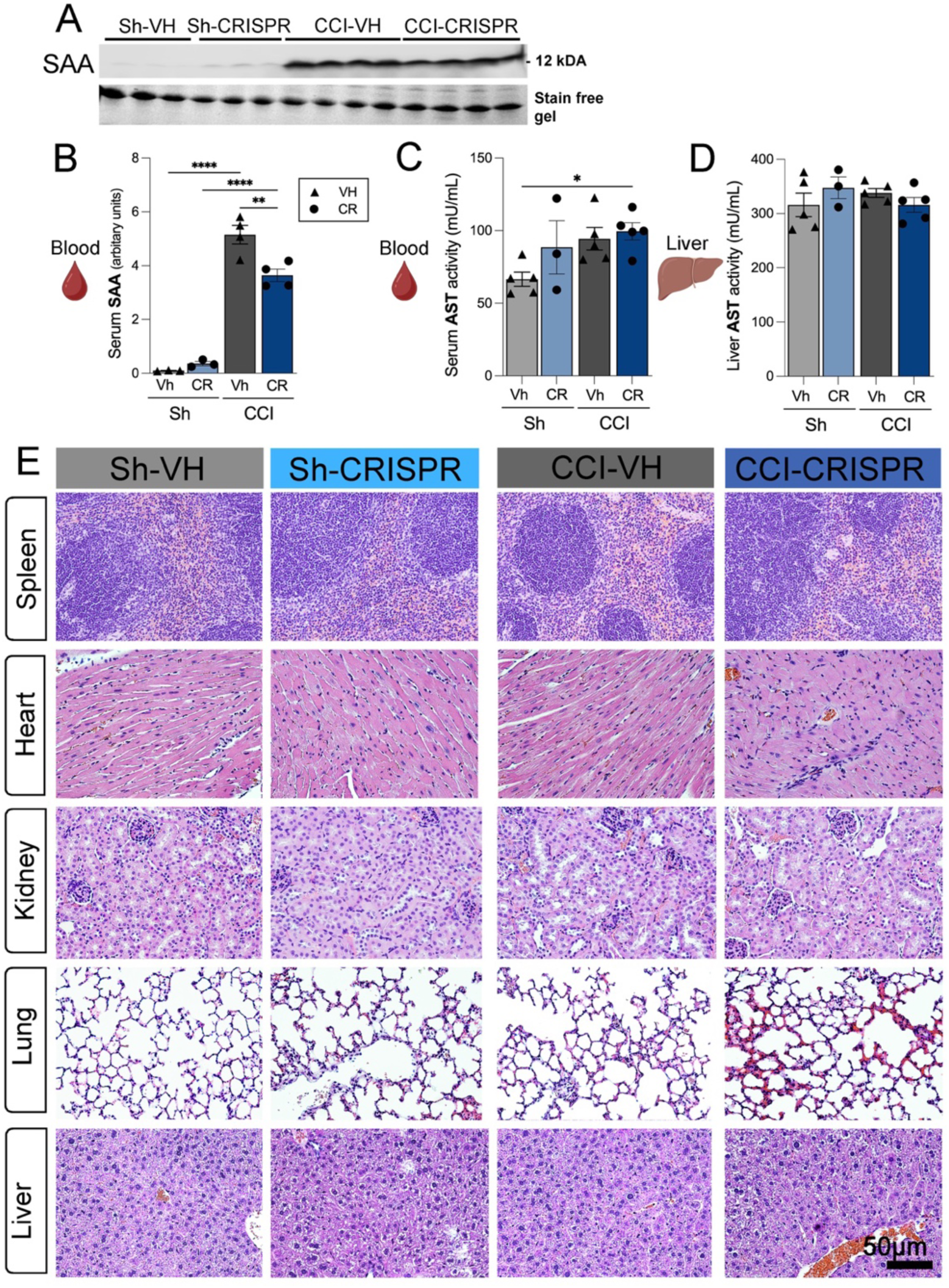
CRISPR-LNP treatment reduces systemic inflammatory markers without inducing peripheral organ toxicity after TBI. (**A**) Representative Western blot of serum amyloid A (SAA) expression in sham (Sh) and controlled cortical impact (CCI) mice treated with vehicle (VH) or CRISPR-LNPs (CRISPR). A stain-free gel is shown as a loading control. **(B)** Quantification of serum SAA levels (arbitrary units), showing a significant increase after CCI that is reduced by CRISPR-LNP treatment. (**C**) Serum aspartate aminotransferase (AST) activity (mU/mL) is a marker of systemic and hepatic injury. (**D**) Liver AST activity (mU/mL), indicating no significant hepatotoxicity associated with CRISPR-LNP treatment. (**E**) Representative hematoxylin and eosin (H&E)-stained sections of spleen, heart, kidney, lung, and liver from Sh-VH, Sh-CRISPR, CCI-VH, and CCI-CRISPR groups. No overt histopathological abnormalities are observed in CRISPR-LNP-treated animals compared to controls. Data are presented as mean ± SEM. *p < 0.05, **p < 0.01, ****p < 0.0001.

## 4 Discussion

Inflammation is a critical protective response to tissue injury; however, when dysregulated, it contributes to disease pathology, and emerging biomaterial-based strategies, particularly nanoscale delivery platforms, offer a powerful approach to enhance the efficacy of anti-inflammatory therapies by improving bioavailability, targeting inflammatory sites, and reducing systemic side effects (Tu et al., 2022). In the present study, we developed and evaluated an Iba-1-targeted CRISPR-LNP platform to modulate microglial polarization through MAPK9 editing in TBI. Our findings demonstrate that: (1) CRISPR-LNPs-mediated MAPK9 targeting promotes a sustainable shift from a pro-inflammatory M1 phenotype to an anti-inflammatory M2 phenotype in macrophages; (2) intranasal delivery significantly enhances preferential targeting of Iba-1+ cells within the brain compared to systemic administration; (3) MAPK9 expression is effectively reduced in the injured cortex *in vivo* after CRISPR-LNP treatment; and (4) this intervention attenuates microglial activation, suppresses pro-inflammatory cytokines, reduces neuronal cell death, and limits peripheral inflammation. Collectively, these results support MAPK9 as a key regulator of neuroinflammation and highlight CRISPR-LNPs-mediated gene editing as a promising therapeutic strategy for TBI.

Neuroinflammation is a central driver of secondary injury following TBI, involving tightly coordinated interactions between central and peripheral immune responses (Boulton & Al-Rubaie, 2025; Donat et al., 2017; Navabi et al., 2024). Microglia, the resident immune cells of the CNS, play a pivotal role in this process. Upon injury, microglia rapidly transition toward a pro-inflammatory phenotype, releasing cytokines and chemokines that amplify immune cell recruitment and exacerbate tissue damage (Kara et al., 2025; Wang et al., 2020; Wei et al., 2021). Although this response is initially protective, its persistence contributes to chronic neurodegeneration. Increasing evidence indicates that shifting microglial polarization toward an anti-inflammatory, reparative phenotype represents a more effective therapeutic strategy than global immune suppression or depletion of microglia, which can impair recovery and worsen outcomes (Shi et al., 2022; Sun et al., 2022; Yang et al., 2019).

MAPK9 (JNK2) is a key component of the stress-activated MAP kinase pathway and has been implicated in inflammatory signaling, apoptosis, and neurodegeneration. By targeting MAPK9 using CRISPR-Cas12a/Cpf1 encapsulated in LNP (Leonard et al., 2020), we achieved a robust suppression of pro-inflammatory gene expression and promotion of M2-associated pathways, even under strong inflammatory stimulation. The observed reduction in chemokines (MCP-1, MIP-1α/β, CCL5, CXCL1) and cytokines (IL-1β, TNF-α) suggests that MAPK9 acts upstream of multiple inflammatory cascades. Importantly, this reprogramming effect was accompanied by morphological changes in microglia, with a reduction in hypertrophic and amoeboid phenotypes, further supporting a functional shift toward a less inflammatory state.

A growing body of evidence indicates that therapeutic strategies promoting a shift from pro-inflammatory M1 to reparative M2 microglial/macrophage phenotypes can significantly improve outcomes. For instance, suppression of SOCS3 expression biases immune cells toward an M2-like state, reducing pro-inflammatory cytokines while enhancing anti-inflammatory signaling and neuroprotection. Similarly, inhibition of TREM-1, a key amplifier of innate immune responses, attenuates neuroinflammation through modulation of the HMGB1-TREM-1 axis and downstream PKCδ/CARD9 pathways, ultimately improving neurological function (Lu et al., 2021). In this context, our findings extend these concepts by demonstrating that targeted CRISPR delivery of MAPK9 using Iba-1-conjugated LNP via intranasal administration specifically enriches these LNP in microglial/macrophage population, and effectively reprograms cell responses toward an anti-inflammatory phenotype. This approach not only suppresses pro-inflammatory mediators but also promotes a permissive environment for tissue repair, aligning with prior studies that highlight the importance of precise immune modulation. Emerging evidence highlights the central role of damage-associated molecular patterns such as HMGB1 in initiating microglial activation through receptors including TLR2, TLR4, and RAGE, leading to NF-κB-dependent inflammatory cascades (Zhang et al., 2022). Targeting these pathways, either pharmacologically or through CRISPR-based gene editing, has been shown to attenuate microglial activation and neuronal injury, underscoring the therapeutic potential of modulating innate immune signaling (Zhang et al., 2022). Accumulating evidence supports the concept that targeting neuroinflammation and enhancing neuroprotective pathways represent a promising therapeutic strategy across a wide spectrum of neurological disorders, including stroke, hypoxia-ischemia (Li et al., 2019; Xie et al., 2021), and TBI. At the molecular level, emerging gene-editing strategies using CRISPR/Cas systems have provided compelling proof-of-concept for directly targeting pathogenic pathways in the brain, including reduction of neuroinflammation and protein aggregation in neurodegenerative diseases. However, a major translational barrier remains the efficient delivery of therapeutics across the BBB. In this context, intranasal delivery and nanoparticle-based platforms have gained considerable attention as non-invasive approaches that enable direct nose-to-brain transport, improving CNS bioavailability. Together, these studies converge on a unifying principle: combining targeted molecular interventions with advanced delivery systems, particularly gene editing and nanotechnology, to mitigate neuroinflammation and improve functional outcomes following brain injury.

The progression of TBI is highly dynamic and influenced by age-dependent and cell-specific glial responses, underscoring the complexity of neuroinflammatory and neurorepair processes. Recent single-cell transcriptomic analyses reveal a diverse landscape of glial subtypes that respond distinctly to injury across developmental stages, highlighting critical signaling interactions such as Notch3-dependent astrocyte responsiveness and CXCL12-CXCR4-mediated microglial recruitment (Qin et al., 2025). In parallel, molecular regulators of gliosis, such as glia maturation factor beta (GMFB), have been shown to drive astrocytic and microglial reactivity, with genetic disruption attenuating reactive gliosis and inflammation following TBI (Yin et al., 2018). Beyond glial activation, excitotoxic mechanisms also contribute substantially to neuronal damage, as evidenced by the role of glutamate carboxypeptidase II (GCPII) in regulating NAAG metabolism and glutamate levels (Ji et al., 2023). Notably, GCPII deletion reduces excitotoxicity and improves cognitive outcomes, whereas its homolog GCPIII appears to play a minimal compensatory role, reinforcing GCPII as a key therapeutic target. Recent advances in gene editing provide a strategy to enhance stem cell resilience, as exemplified by CRISPR/Cas9-mediated disruption of Keap1 in mesenchymal stem cells, which activates the Nrf2 antioxidant pathway, reduces oxidative damage, and promotes cell survival through modulation of apoptotic and proliferative signaling. These findings highlight the importance of reinforcing intrinsic cellular defense mechanisms to improve therapeutic engraftment.

Another key innovation of this study is the use of LNPs for CRISPR delivery. LNPs represent the most clinically advanced non-viral gene delivery platform (Hosseini-Kharat et al., 2025), with demonstrated safety and scalability (Hoy, 2018; Lamb, 2021). Our formulation, based on ionizable lipids (Kulkarni et al., 2018), enables efficient encapsulation of CRISPR components while minimizing immunogenicity. Utilization of LNP also allows for surface functionalization with Iba-1 antibodies, which further enhances cell-type specificity to allow preferential targeting of microglia/macrophages (Henriksen-Lacey et al., 2011). This targeted approach is particularly important in the CNS, where non-specific gene editing could lead to unintended effects in neurons or other glial populations. In our study, Iba-1-targeted CRISPR-LNPs were uptaken selectively by Iba-1+ microglia cells after intranasal administration, with minimal uptake by Iba-1- cell populations. Moreover, CRISPR-LNPs targeting MAPK9 rapidly induced double-strand DNA breaks in macrophages within minutes of internalization (Horikoshi et al., 2019), with a significant reduction in cortical MAPK9 mRNA levels in mice at 1dpi. *In vitro*, MAPK9 editing suppressed M1 polarization while preserving an M2-like phenotype, as reflected by an increased proportion of M2-like cells and a reduction in M1-like cells. Delivery across the BBB remains a major challenge for CNS therapeutics. Intranasal administration has gained attention as an effective, non-invasive route for CNS drug delivery, enabling direct nose-to-brain transport while bypassing the BBB. Here, we demonstrate that intranasal administration provides a highly effective, non-invasive route for delivering CRISPR-LNPs to the brain. Compared to intravenous administration, intranasal delivery significantly increased uptake in Iba-1+ cells while reducing off-target distribution. This likely reflects transport along olfactory and trigeminal pathways, as well as perivascular and lymphatic routes, enabling rapid access to the brain parenchyma. These findings reinforce the translational potential of intranasal delivery for gene therapies targeting neuroinflammation. Functionally, MAPK9 editing resulted in a rapid attenuation of neuroinflammatory responses following TBI. We previously demonstrated that serum SAA is an acute-phase protein produced in the liver after TBI (Villapol et al., 2015; Wicker et al., 2019). Beyond central effects, CRISPR-LNP treatment also reduced systemic inflammation, as evidenced by decreased serum SAA levels. This finding underscores the bidirectional interaction between brain injury and peripheral immune responses and suggests that targeting central inflammatory pathways can have broader systemic benefits. Notably, safety analyses revealed no evidence of peripheral toxicity, as indicated by stable AST levels and the absence of histopathological abnormalities in major organs. These results support the favorable safety profile of this approach.

This study has several limitations. First, the optimal dosing and timing of CRISPR-LNP administration remain to be defined. Second, although MAPK9 targeting produced robust anti-inflammatory effects, the precise downstream signaling mechanisms require further investigation. Third, our analyses were limited to the acute phase, and future studies should evaluate long-term outcomes, including cognitive and functional recovery. Finally, differences between murine and human microglial biology warrant validation in human-relevant systems before clinical translation.

In conclusion, this study establishes a novel and targeted therapeutic framework for TBI by integrating gene editing, nanomedicine, and non-invasive delivery to directly modulate neuroinflammatory signaling at its source. Unlike conventional anti-inflammatory approaches, our strategy leverages intranasal CRISPR-LNP technology to selectively disrupt MAPK9-dependent pathways within microglia, thereby reprogramming the inflammatory response from a chronic, neurotoxic state toward a reparative phenotype during the peak of inflammation in TBI (Villapol et al., 2014; Villapol et al., 2017). This work not only identifies MAPK9 as a key regulator of sustained neuroinflammation but also demonstrates the feasibility and efficacy of precision gene editing in the injured brain. Importantly, the combination of cell-specific targeting and nose-to-brain delivery overcomes major translational barriers, offering a scalable and clinically relevant platform. Collectively, our findings provide a proof-of-concept for next-generation neurotherapeutics that move beyond symptom management to directly rewire pathological processes, opening new avenues for the treatment of TBI and other neuroinflammatory disorders.

## Declarations

Competing interests The authors declare no competing interests.

## Author Contributions

F.L. and S.V. conceived and designed the study. G.K. led the experimental work, performed data analysis, and drafted the manuscript. F.L. and T.G. prepared the nanoparticles. Y.A., J.L.-E., P.P., M.H., H.F., and N.T. contributed to methodology development, data acquisition, and analysis, and critical revision of the manuscript. S.V. supervised the project, secured funding, and contributed to study design and manuscript editing. All authors reviewed and approved the final manuscript.

